# Hepatitis C virus replication fitness as a determinant of antiviral therapy outcome

**DOI:** 10.64898/2026.02.03.703674

**Authors:** Ha Gyu-Thomas Seong, Tomke Arand, Colin Förster, Micha Böhm, Eva Heger, John McLauchlan, Christoph Sarrazin, Julia Dietz, Rolf Kaiser, Volker Lohmann, Paul Rothhaar

**Author notes:** Co-corresponding authors. Volker Lohmann, Paul Rothhaar. These authors contributed equally (Ha Gyu-Thomas Seong, Tomke Arand).

## Abstract

**Background and Aims:** Hepatitis C virus (HCV) infections were previously treated with interferon (IFN) but today direct acting antivirals (DAAs) with cure rates >95% are available. DAA treatment failure is primarily attributed to resistance associated mutations (RAMs), often imposing a fitness cost. Interferon treatment outcome was shown to be associated with the interferon sensitivity determining region (ISDR), which is part of the replication enhancing domain (ReED) in non-structural protein (NS) 5A. We found that accumulation of mutations in the ReED was indicative of elevated viral genome replication fitness. This study investigates the impact of HCV replication fitness on antiviral treatment outcomes.

**Methods:** We utilized chimeric HCV subgenomic replicons containing RAMs and ReED sequences from patients after interferon treatment or DAA failure to assess replication fitness of patient isolates in presence and absence of inhibitors.

**Results:** Replication fitness did not impact on IFN sensitivity in cell culture but resulted in higher remaining antigen levels for highly replicating variants at a given IFN concentration. Furthermore, we identified ReED variants substantially increasing HCV replication in several patients who failed DAA therapy across different genotypes. High replicator ReEDs rescued the fitness loss caused by RAMs like Y93C/H (NS5A) and S282T (NS5B). While high replication fitness did not intrinsically increase drug sensitivity (IC50), it allowed the virus to sustain robust replication despite antiviral pressure.

**Conclusions:** Elevated replication fitness might support interferon treatment due to increased antigen presentation, facilitating adaptive immune responses. Furthermore, ReED mediated increase in replication fitness could contribute to DAA treatment failure by preserving higher replication upon treatment and compensating for RAM associated fitness costs. Thus, patients failing DAA treatment should be monitored for RAMs and ReED mutations.

## Introduction

Hepatitis C virus (HCV) infections persist in ∼75% of patients [1] remaining a public health issue with 50 million people infected and 240,000 dying from the disease in 2022 [2]. Current therapy regimens based on direct acting antivirals (DAAs) offer cure rates >95% (reviewed in [3]). Beforehand, treatment strategies relied on pegylated interferon alpha 2a (IFN) in combination with ribavirin achieving cure rates of ∼50% [4]. IFN acts as a cytokine which induces the production of antiviral interferon stimulated genes and potentially has other immunomodulatory functions [5]. HCV is a heterogenous virus classified into 8 genotypes with over 100 subtypes [6]. This heterogeneity is enabled by the error-prone genome replication cycle of HCV also leading to the emergence of a variety of closely related viral variants within each patient, the so-called quasispecies [7]. HCV has a positive sense RNA genome coding for one open reading frame. The resulting polyprotein is cleaved into individual viral proteins by host and viral proteases. The non-structural proteins (NS) 3 to NS5B form the viral replicase which together with the untranslated regions (UTRs) flanking the viral genome are sufficient for HCV genome replication (reviewed in [8]). NS3 has helicase activity and can function as a protease together with the co-factor NS4A. NS4B is needed for membrane rearrangements to form the viral replication organelle. NS5A is a phosphoprotein without enzymatic activity that interacts with a plethora of host and viral proteins. NS5B is the RNA dependent RNA polymerase needed for replication of the RNA genome. There are three classes of DAAs in clinical use targeting distinct HCV proteins: NS3-protease inhibitors like Telaprevir, NS5A inhibitors like Daclatasvir (DCV) and the nucleotide analogue Sofosbuvir (SOF) targeting NS5B (reviewed in [3]). A major reason for treatment failure is based on the virus’ ability to rapidly adapt to the treatment by acquiring resistance associated mutations (RAMs) within the viral genome [9]. For DCV, the RAMs cluster in domain 1 of NS5A with L31M or Y93H being prominent examples [10]. For SOF, the RAM S282T was identified although it is rarely detected in vivo, potentially due to inducing a reduction in replication fitness [11, 12]. To combat RAM induced treatment failure, DAAs are usually given as a combination therapy of at least two drug classes (NS3 + NS5A inhibitor or NS5A + NS5B inhibitor) [13].

DAAs were developed based on the subgenomic replicon (SGR) system allowing for HCV studies in cell culture [14]. Here, the HCV replicase (NS3-5B and the UTRs) is expressed together with a reporter or resistance protein. Initially, the virus needed to acquire cell culture adaptive mutations for efficient replication [15] but improvements to the cell culture system like overexpression of the cytosolic lipid transported SEC14L2 allowed for studies on the replication of wildtype HCV constructs [16]. Based on reporter SGRs, we previously identified the replication enhancing domain (ReED) within NS5A, where we found that mutations caused an up to 1000-fold increase in genome replication fitness (RF) [17]. The sequence signature of elevated RF was an accumulation of amino acid mutations in the N-terminal half of the ReED [17], in a region previously associated with interferon treatment, the interferon sensitivity determining region (ISDR), for which accumulation of four or more mutations were found to be associated with efficient response to IFN therapy [18]. In the following, we investigated the interplay of ReED mediated replication enhancement, RAMs and antiviral treatment response in order to assess whether sequencing the ReED of HCV infected patients might help in understanding reasons for treatment failure. Thus, we assessed the relevance of the ReED in immunocompetent patients based on an IFN treatment cohort. Additionally, we identified and studied patients who failed DAA treatment and harboured a ReED allowing for elevated RF. We further studied the fitness loss induced by some RAMs and the subsequent rescue by a high replicator ReED as well as the response of high replicators containing RAMs to DAA treatment in cell culture.

## Results

### ISDR mutations correlate with high replication fitness in patients of an IFN treatment cohort

Treatment regimens based on IFN were the most common strategy to cure HCV infected patients until the establishment of DAAs in the 2010s [3]. In 1995, a correlation between accumulation of mutations in the amino acid sequence of the ISDR with the response to IFN treatment was observed for gt1b [18]. The ISDR is a subregion of the ReED which is a regulator of RF located within the NS5A protein of HCV (Fig. 1A, upper panel) [17]. Interestingly, we previously identified 3 or more ISDR mutations of a given isolate when compared to the gt1b consensus as a sequence signature of elevated RF in liver transplant patients [17]. So far, we did not characterise ReEDs with altered ISDRs from patients outside the transplant context. Therefore we studied the ReEDs of 4 IFN resistant and 11 IFN sensitive gt1b patients from the initial ISDR cohort (Fig. 1A, lower panel) [18] for replication in cell culture, using SGRs based on the HCV prototype isolate Con1 [14], harbouring the respective patient derived ReEDs (Fig. 1B). 5 out of 7 ReEDs with 3 or more ISDR mutations, all derived from IFN sensitive patients, showed an RF more than 100 -fold above the replication level of a control construct containing the gt1b consensus ReED (Fig. 1C). In contrast, none of the ReEDs with less than 3 ISDR mutations, including all from IFN resistant patients, showed such a high replicator phenotype. Beyond gt1b, two ReEDs with 8 and 4 ISDR mutations respectively from gt1a infected patients which were successfully treated with IFN [19] elevated RF of the gt1a prototype isolate H77 (Fig. S1A). This confirmed the connection between 3 or more ISDR mutations and elevated RF in immunocompetent patients. Nevertheless, with P12_IFN(s) and P16_IFN(s), we identified two gt1b patients who were IFN sensitive and harbored an isolate with 5 or 6 ISDR mutations, respectively, without showing a high replicator phenotype in cell culture (Fig. 1C). To get a deeper understanding of these isolates, we only focused on the ISDR and replaced the C-terminus (C-term) of the ReED of the patient isolates with the gt1b consensus sequence. While the construct with the P16_IFN(s) ISDR remained a low replicator, the P12_IFN(s) ISDR increased RF nearly 100-fold showing that P12_IFN(s) had a high replicator ISDR with a C-term downregulating RF to a low replicator level (Fig. S1B). This confirmed previous results of the ISDR being the main driver of elevated RF [17].

**Figure 1:**
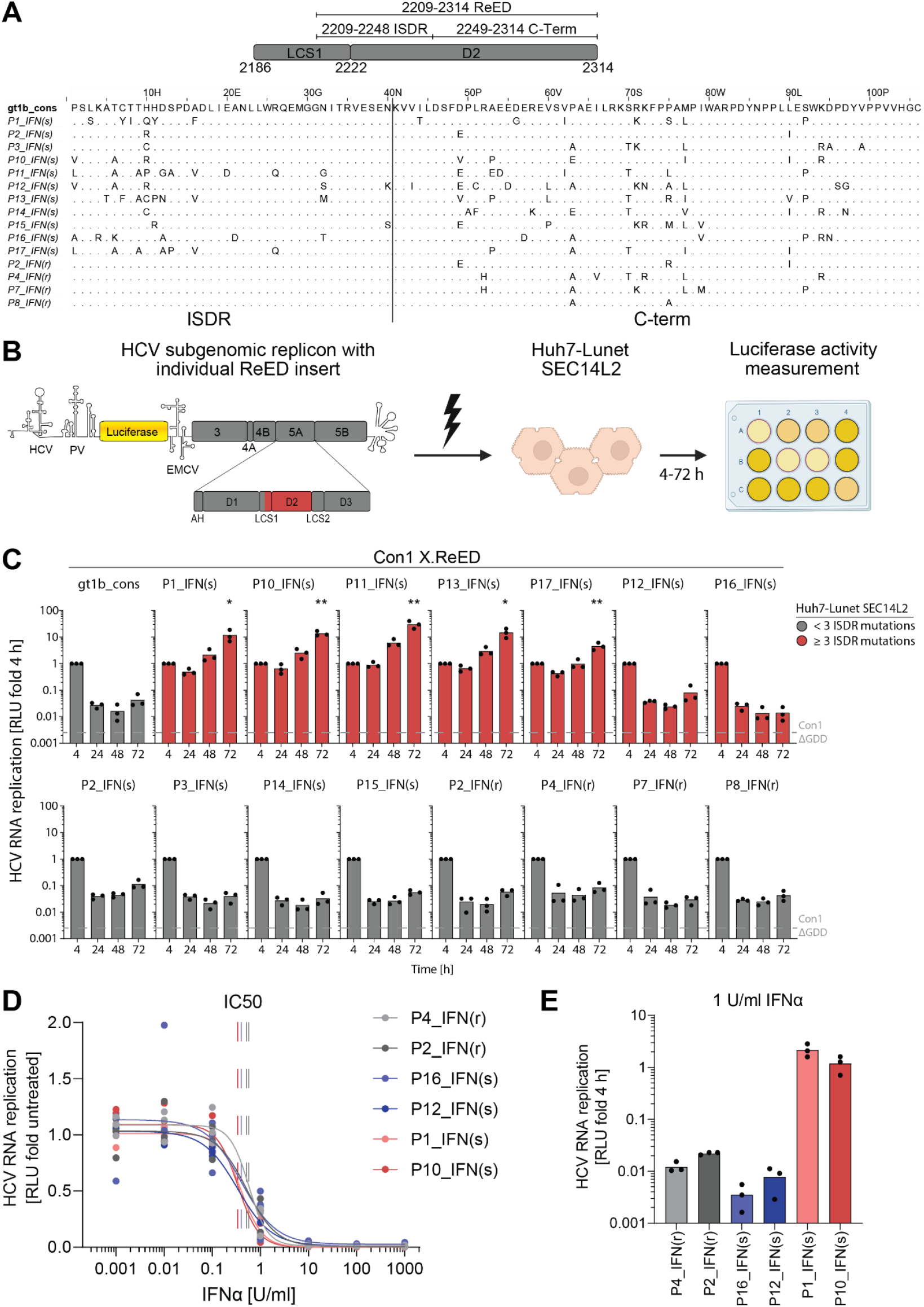
HCV replication fitness and IFN treatment. A) Schematic of the position of ReED, ISDR and C-term within low complexity sequence 1 (LCS1) and domain 2 (D2) of NS5A. Position numbering based on the Con1 polyprotein sequence (upper panel). Alignment of the ReED amino acid sequences of all isolates from the IFN treatment cohort [18] with a genotype 1b consensus ReED serving as a reference. Isolate nomenclature was taken from [18]. Dots indicate that an amino acid is identical to the consensus (lower panel). B) Schematic of the experimental setup. Con1 subgenomic replicons harbouring a patient derived ReED were transfected via electroporation into Huh7-Lunet SEC14L2 cells. A luciferase reporter served as readout for replication fitness. C) Con1 SGRs harboring the indicated ReED were electroporated into Huh7-Lunet SEC14L2 cells, luciferase activity in cell lysates (RLU) was quantified as a correlate of RNA replication efficiency at the given time points and normalized to 4 h to account for differences in transfection efficiency. The dashed grey line indicates the 72 h time point for Con1 ΔGDD, a replication deficient negative control. Data are from three independent biological replicates measured in technical duplicates. Each dot depicts the result of one replicate and the bar indicates the mean of all replicates. D) Huh7-Lunet SEC14L2 cells electroporated with Con1 SGRs harboring the indicated ReED were treated with variable IFN concentrations 24 h after electroporation. Depicted is the replication 72 h after treatment of Con1 SGRs with ReEDs from either IFN resistant patients (P2_IFN(r)&P4_IFN(r)) or IFN sensitive patients with a low replicator (P12_IFN(s)&P16_IFN(s)) or a high replicator ReED (P1_IFN(s)&P10_IFN(s)) normalized to an untreated control. Data are from three independent biological replicates measured in technical duplicates with each dot depicting the result of one replicate. IC50 was determined through a nonlinear fit shown as a continuous line, the dashed line marks the IC50. E) Data from panel D showing the replication 72 h after treatment with 1 U/ml IFN normalized to the input 4 h after electroporation. C) Statistical significance between the 72 h time point of the indicated isolates and the 72 h time point of the Con1 gt1b_cons.ReED sample was determined using a two-sided Student’s t-test. * = p < 0.05, ** = p < 0.01.

The results from the IFN treatment cohort (Fig. 1C) further implied that patients harbouring an HCV high replicator variant apparently had a better response to IFN treatment, suggesting that high RF might directly increase IFN sensitivity. However, exogenous treatment with IFN showed similar relative inhibitory effects on low and high replicators, with IC50 values ranging from 0.2-0.6 U/ml (Fig. 1D, S1C), irrespective of treatment outcome in vivo. Conversely, when looking at the absolute replication levels in presence of 1 IU/ml IFN, a concentration slightly above the IC50, high replicators still showed 100-fold higher levels of luciferase activity when compared to low replicators (Fig. 1E, S1D). In patients, the increased replication of a high replicator in presence of IFN therefore might lead to substantially higher antigen presentation subsequently facilitating the detection of infected cells by the adaptive immune response. Overall, the potential connection of high RF and a patient’s IFN sensitivity might not be caused by intrinsic changes of the viral IFN sensitivity but might rather become apparent in the interplay of the virus with a fully functional immune system. High RF thereby appears to contribute but is not a prerequisite of IFN treatment response, since several low replicating variants were found in patients successfully cured by IFN therapy.

### DAA treatment failure patients can harbour a high replicator HCV variant sometimes emerging over the course of antiviral therapy

DAA treatment only fails in a small fraction of patients and is often attributed to the virus acquiring RAMs which reduce the viral sensitivity to the DAAs. We wanted to assess whether mutations enhancing viral replication fitness might be a so far overlooked factor affecting DAA treatment responses. Again, we used 3 or more amino acid mutations in the ISDR (Fig. 1A) as the sequence signature of high HCV genome replication fitness and analysed the frequency of such potential high replicators in the context of DAA treatment failure. In a dataset of a total of 971 patients combining results from HCV Research UK [20], PEPSI [21] and the European DAA resistance database at the University Hospital Frankfurt, we found no enrichment of patients infected with a potential HCV high replicator variant in DAA failure cases (Fig. 2). Nevertheless, while not enriched we still identified 9 patients from all three tested gts 1a, 1b and 3a that failed DAA treatment and contained a potential high replicator variant. We tested all of those for high replicator phenotypes in a genotype matched replicon backbone.

**Figure 2:**
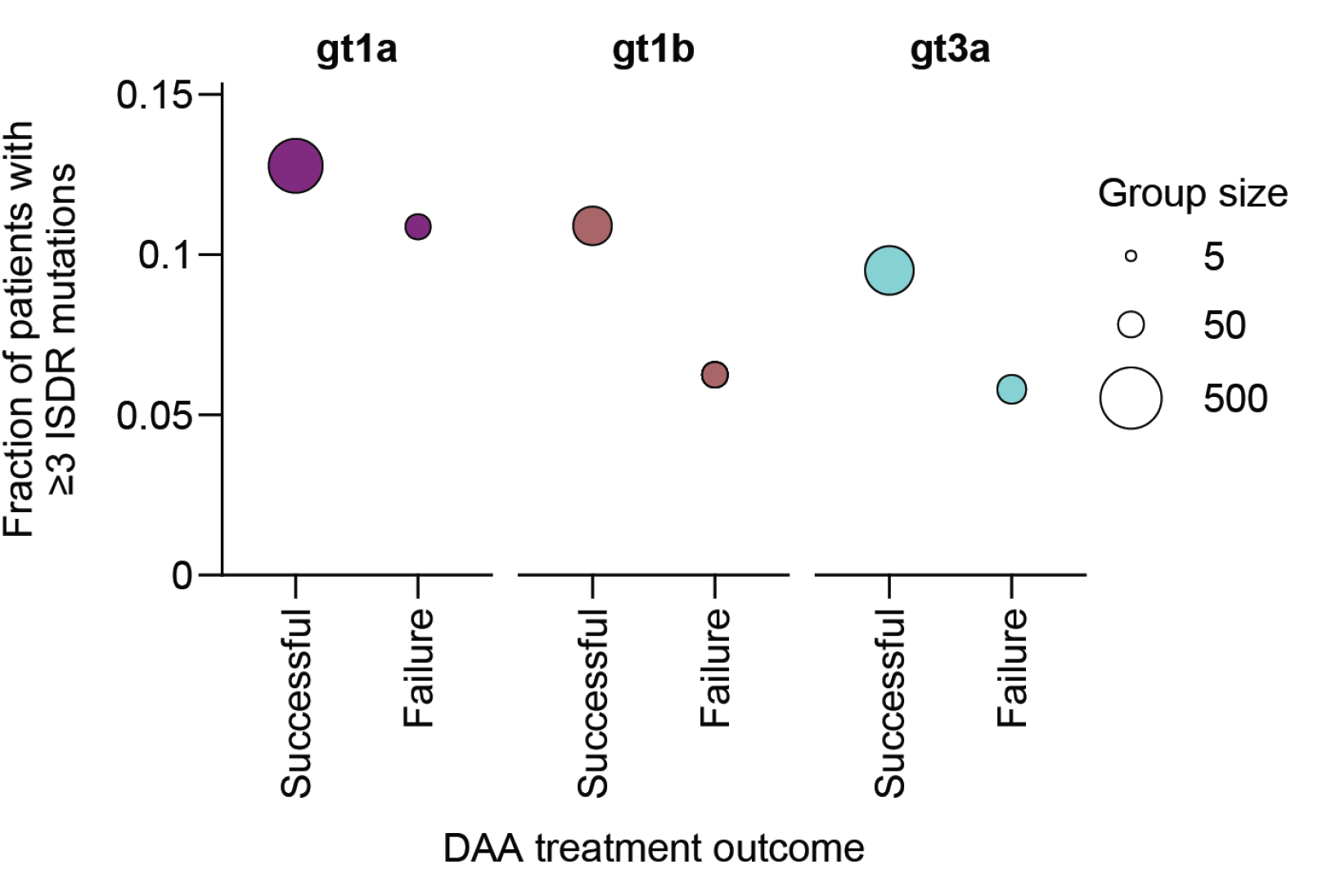
Frequency of potential high replicators in the context of DAA treatment response. For each patient’s ISDR sequence, the number of amino acid differences compared to the gt specific consensus sequence was determined. If an ISDR had 3 or more mutations, the patient was considered to be infected with a potential high replicator. Data based on a cohort combining patients from HCV Research UK [20], PEPSI [21] and the European DAA resistance database at the University Hospital Frankfurt.

For gt1a, the ReED sequences of 4 patients were introduced into the SGR backbone of isolate H77 [17], employing a similar experimental setup as for RF assessment in the IFN cohort (Fig. 1B). Indeed, 3 out of 4 tested gt1a ReEDs enhanced RF by ∼10-fold when compared to a reference low replicator construct harbouring the gt1a consensus ReED (Fig. 3A). Analogous, we tested 2 ReEDs from gt1b patients in the prototype Con1 SGR with isolate DAA_6 increasing replication 7-fold compared to the gt1b consensus ReED (Fig. 3B). In case of the gt3a DAA treatment failure patients, isolate S52 was used [17] with all 3 tested ReEDs increasing RF by 10-1000-fold compared to the gt3a consensus control (Fig. 3C). Therefore, 7 out of 9 tested ReEDs with high replicator sequence signatures were verified as phenotypical high replicators.

**Figure 3:**
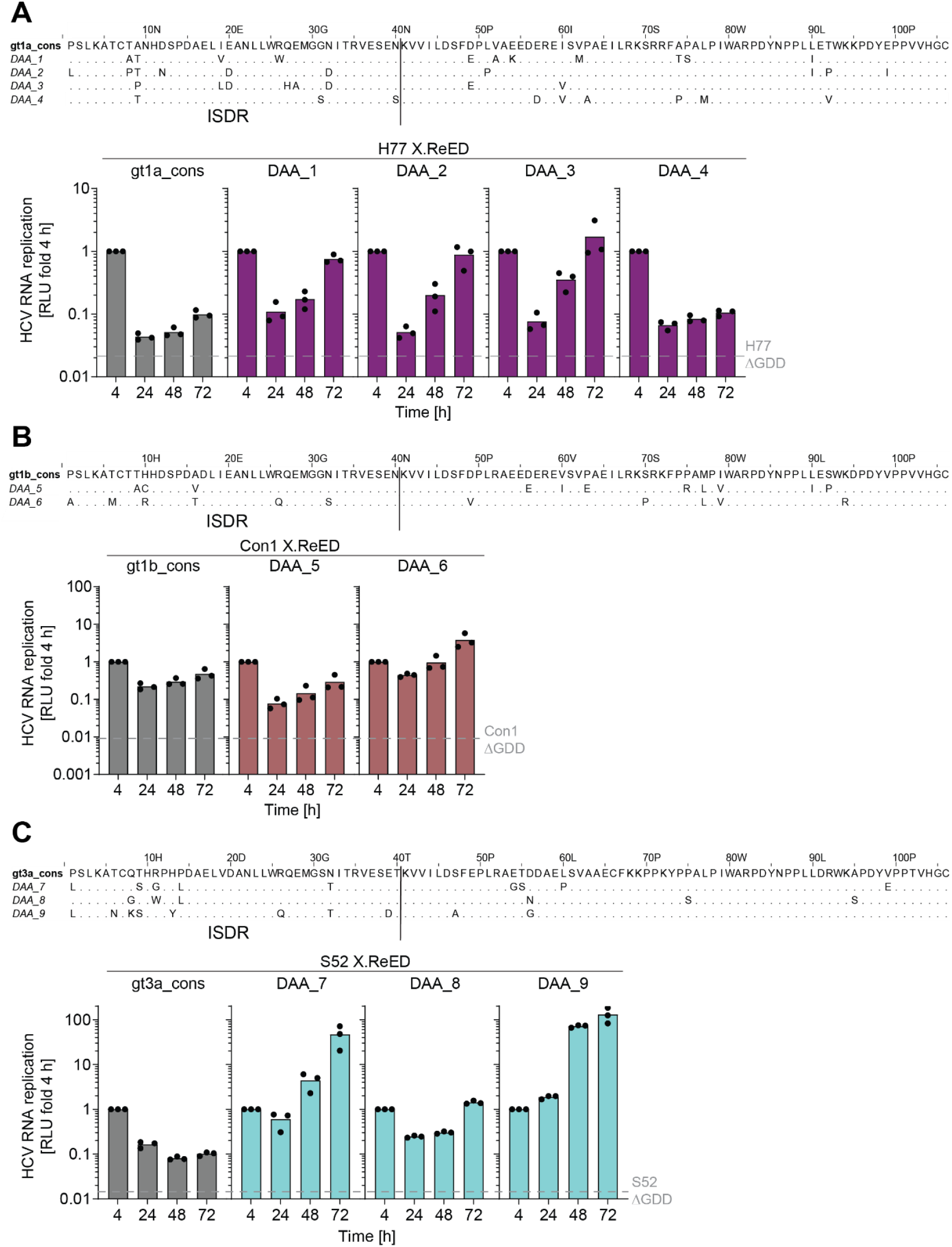
Replication fitness of selected ReEDs from DAA failure patients. A-C) Alignments of the tested ReEDs with the genotype specific amino acid consensus sequence (upper panels). Replication studies were performed as described in Fig. 1B. Chimeric prototype SGRs H77 (A), Con1 (B) or S52 (C) harboring the indicated ReEDs were used. The dashed grey line indicates the 72 h time point for H77/ Con1/ S52 ΔGDD, replication deficient negative controls. Data are from three independent biological replicates measured in technical duplicates. Each dot depicts the result of one replicate and the bar indicates the mean of all replicates.

Serial samples were available for a subset of patients with most of them harbouring a high replicator sequence already prior to treatment initiation, suggesting that the presence of a high replicating variant might have facilitated the emergence of DAA resistance (Fig. 4A). Interestingly, in case of 3a_LTX_2, a high replicator variant identified in a previous study [17], therapy failed after LTX. In this case, immunosuppression might have contributed to the emergence of the high replicator. Only in two patients, high replicating variants evolved during DAA treatment: In patient DAA_8, an increase in RF emerged over the course of the failed DAA treatment. Patient DAA_6 failed DAA treatment twice (Fig. 4B). For this patient, NGS data were available, showing that the high replicator variant was already present as a non-dominant species before treatment initiation but became dominating only after the second treatment failure (Fig. 4C).

**Figure 4:**
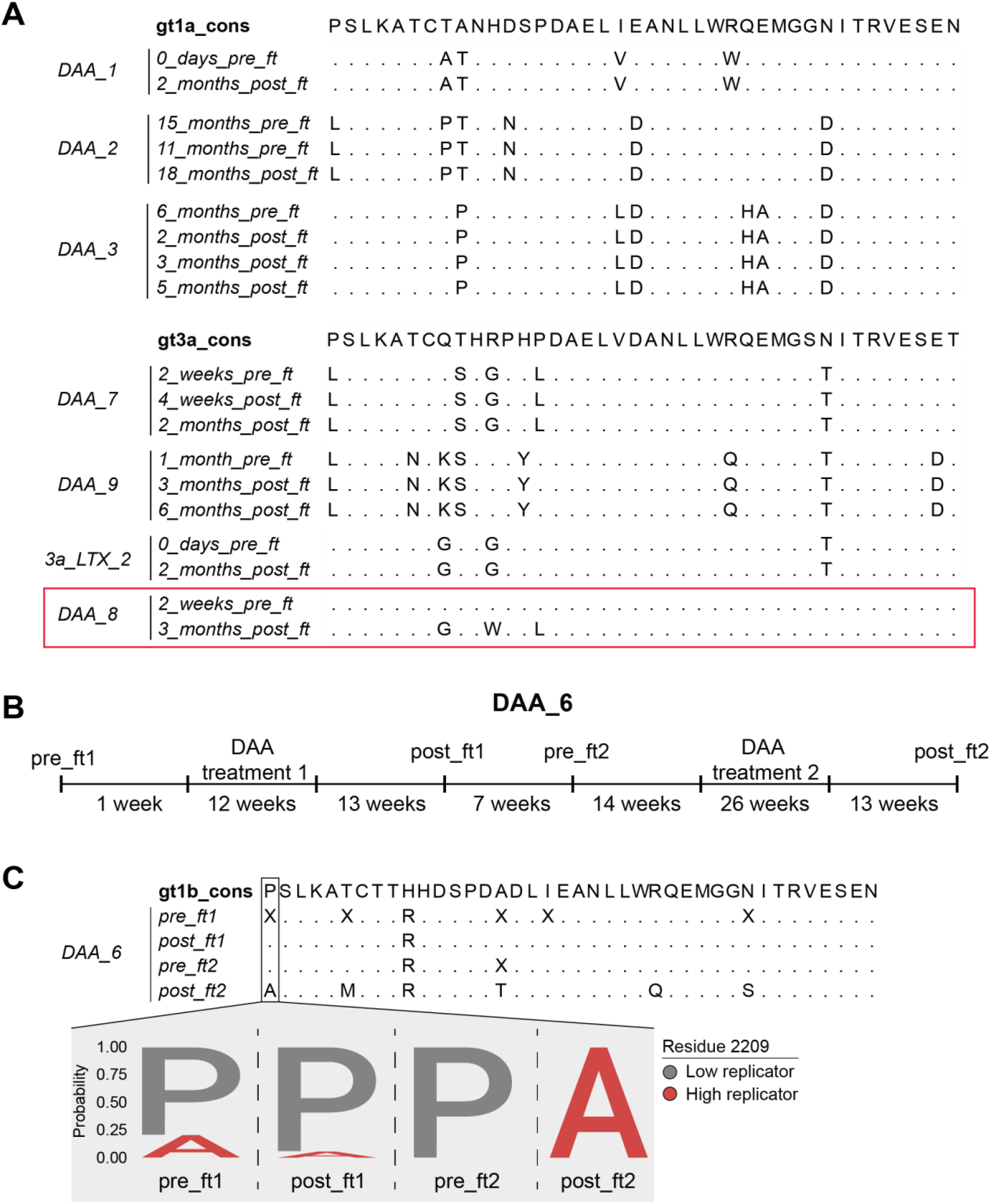
ISDR evolution in patients failing DAA treatment. A) Alignment of ISDR sequences from before and after failed DAA treatment (ft) with the respective genotype specific consensus for all g1a and gt3a patients where this information was available. Note that only in patient DAA_8 a change in the ISDR over the course of DAA treatment was observed (red box). B) Timeline of sample collection and DAA treatment of patient DAA_6. DAA treatment 1 was performed with ribavirin, ledipasvir and SOF, DAA treatment 2 was performed with ribavirin, simeprevir and SOF. The ReED from time point post_ft2 was used in Fig. 3B. C) Alignments of the ISDR consensus sequences from all time points with the gt1b consensus sequence. X indicates a polymorphism (upper panel). Sequence logo based on NGS results representing the relative frequency of proline and alanine at different time points in the DAA_6 patient (lower panel). P2209A was previously shown to enhance RF [17].

These results highlight that, at least in some cases, the quasispecies dominating after failure of DAA therapy had a high replicator phenotype. In most cases, the high replicator variants were already present prior onset of therapy, whereas in two patients a selection of a high replicator over the course of a failed DAA therapy occurred.

### High replicator ReEDs mediate elevated replication fitness in the context of RAMs and DCV treatment

A main reason for DAA treatment failure is the virus acquiring RAMs. Often, these mutations come at a fitness cost [22]. Therefore, we hypothesized that a high replicator ReED could mitigate this fitness cost and thus help HCV to withstand the antiviral therapy. For gt1a, we found the DCV RAM L31M [23] in patient DAA _2 and RAM Y93C [23] in patient DAA _3 which both also had a high replicator ReED (Fig. 3A). Introduction of L31M into the gt1a construct had little impact on RF while Y93C reduced replication by 65% (Fig. 5A). The ReED sequences of DAA_2 and DAA_ 3 both strongly increased RF also in combination with RAMs L31M and Y93C, respectively. Thus, the ability of a high replicator ReED to elevate RF is maintained in the presence of RAMs and in case of Y93C it exceeded the fitness cost of the RAM by far.

**Figure 5:**
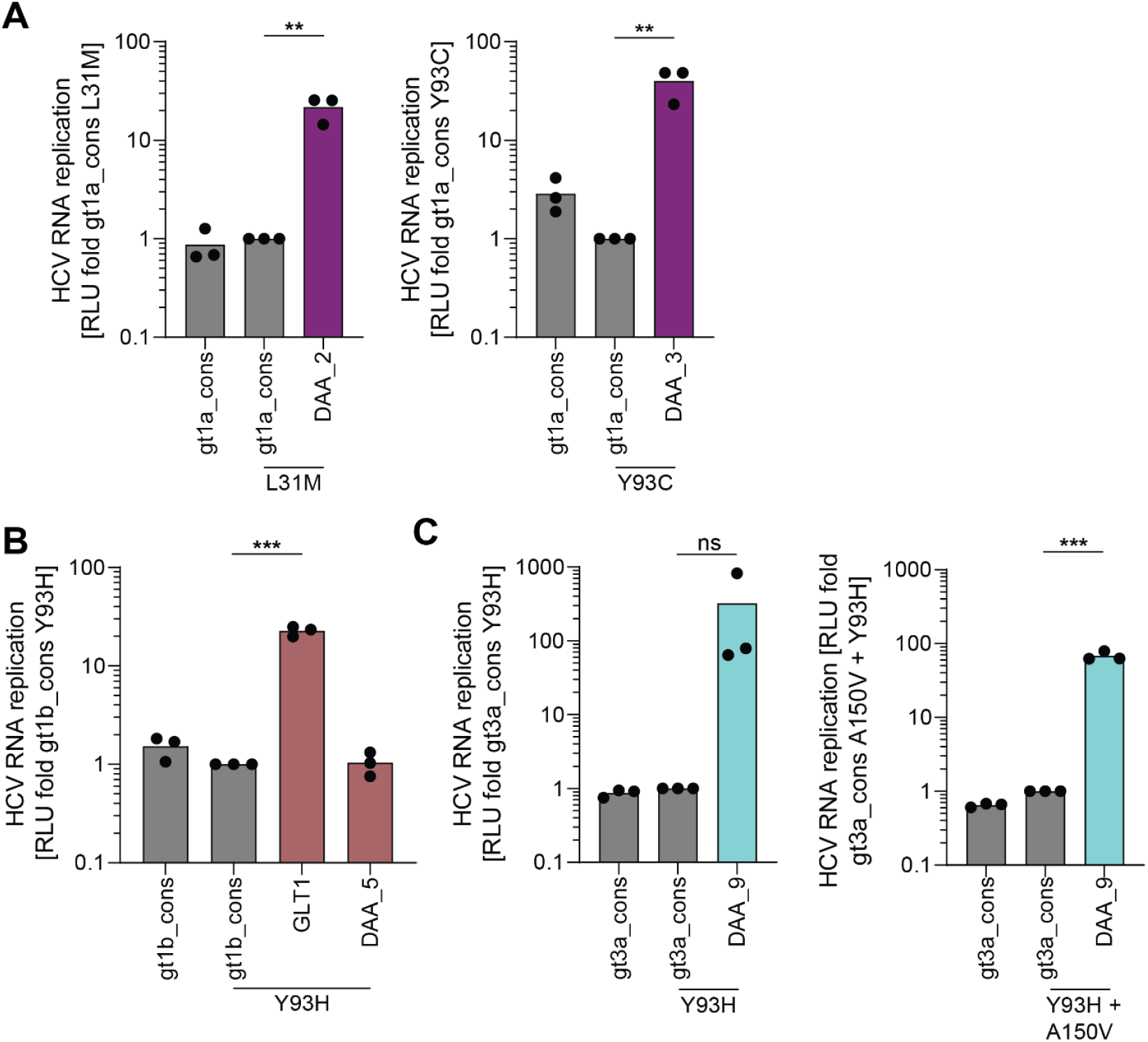
High replicator ReEDs can compensate for the fitness loss induced by some DCV RAMs. A-C) Replication studies were performed as described in Fig. 1B. Results were first normalized to 4 h to account for differences in transfection efficiency and then normalized to the RNA replication of the construct with the gt specific consensus ReED and the indicated RAM. Displayed is the HCV RNA replication 72 h after electroporation. Chimeric prototype SGRs H77 (A), Con1 (B) or S52 (C) harboring the indicated ReEDs and the indicated RAMs were used. Data are from three independent biological replicates measured in technical duplicates. Each dot depicts the result of one replicate and the bar indicates the mean of all replicates. Statistical significance was determined using a two-sided Student’s t-test. ns = not significant (p > 0.05), ** = p < 0.01, *** = p < 0.001.

For gt1b, DAA_ 6 contained the DCV RAM Y93H [23]. Since the DAA_6 ReED had a low replicator phenotype, we further included the bona fide high replicator ReED of the GLT1 isolate [17, 24] to analyse the impact on replication in presence of the RAM. Y93H caused a 30% replication fitness reduction when introduced into a Con1 SGR with the gt1b consensus ReED (Fig. 5B). The ReED of DAA_ 6 did not differ from Con1 with the gt1b consensus ReED, as expected from its low replicator phenotype (Fig. 5B). However, the GLT1 ReED mediated a 22-fold increase in replication fitness in the presence of Y93H showing that, in principle, a high replicator ReED can also mitigate the RAM induced fitness loss in gt1b.

Lastly, we assessed the gt3a patient DAA_9 who harboured both Y93H in NS5A and the SOF RAM A150V [25] in NS5B. Surprisingly, Y93H as well as the combination of Y93H and A150V slightly increased RF of S52 with the gt3a consensus ReED (Fig. 5C). Again, the high replicator ReED from patient DAA_9 strongly boosted RF also in the presence of the RAMs. In conclusion, only some of the RAMs found in our patient cohort reduced RF but, in all cases, characterisation of the RAM + ReED combinations found in individual patients revealed that the presence of a high replicator ReED strongly enhanced RF, compensating for all negative effects on RF induced by the RAMs.

Patients DAA_2&3 (both gt1a) and DAA_9 (gt3a) were all shown to have high replicator ReEDs, DCV specific RAMs and failure of DAA treatment with regimens containing DCV. Upon treatment with DCV at its IC90 of 0.09 nM (gt1a) and 1.3 nM (gt3a) (Fig. S2), replication of all constructs was reduced but those combining a RAM with a high replicator ReED had a significant advantage with 10- to 100-fold higher RF than constructs with a low replicator ReED (Fig. 6A&C). This replicative advantage upon DCV treatment was also maintained in the absence of RAMs, when testing previously published high replicator ReEDs from post LTX patients (Fig. 6B&D) [17]. The results imply that a high replicator ReED might primarily act as a fitness buffer, facilitating the emergence of RAMs under selective pressure of DAA therapy and/or mitigating the potential fitness cost associated with RAMs.

**Figure 6:**
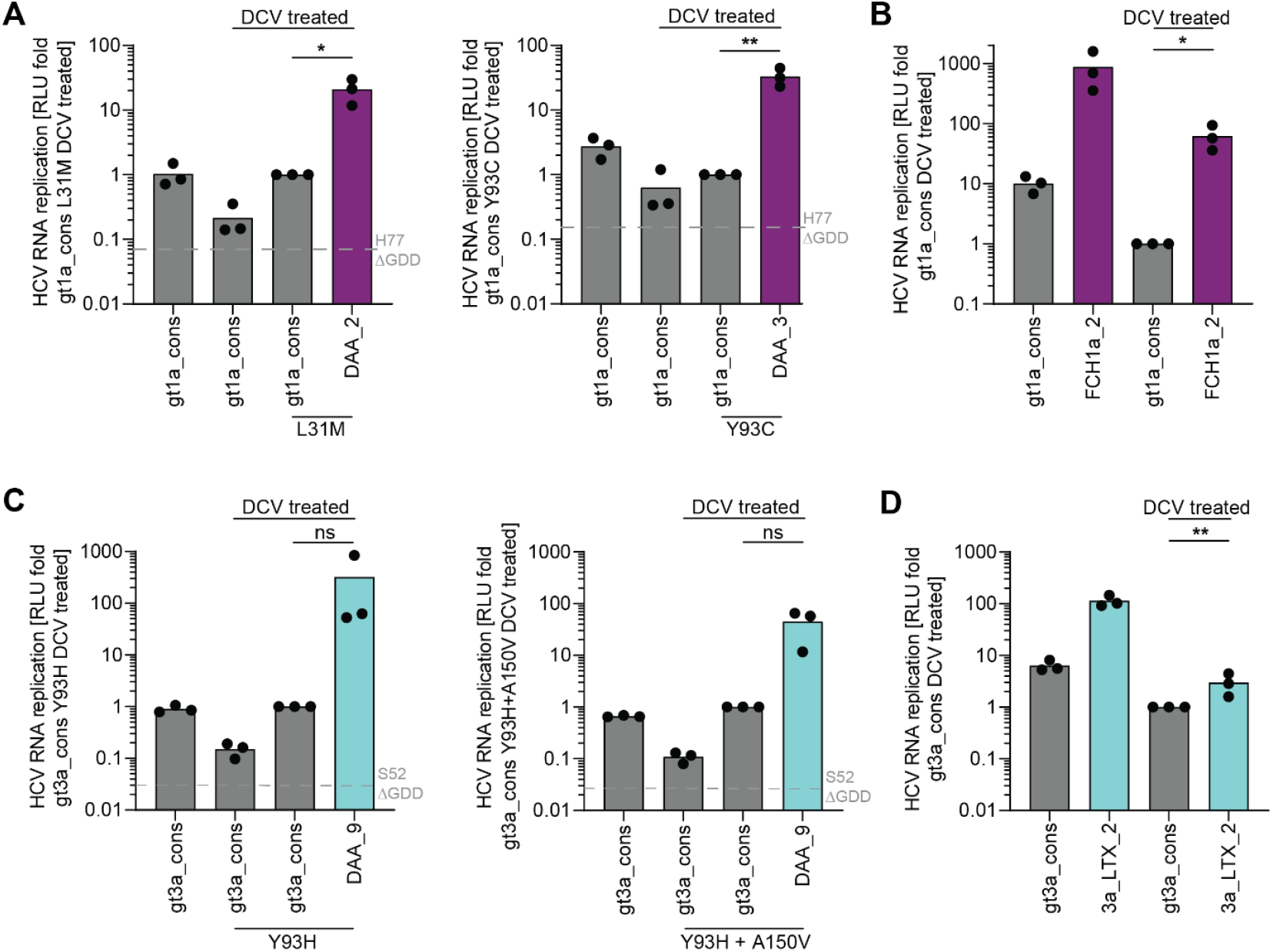
High replicators maintain elevated replication levels upon DCV treatment. Replication studies were performed as described in Fig. 1B. Results were first normalized to 4 h to account for differences in transfection efficiency and then normalized to the RNA replication of the DCV treated construct with the gt specific consensus ReED and the indicated RAM. DCV treatment with the IC90 concentration (gt1a 0.09 nM, gt3a 1.3 nM) as determined in Fig. S2 was performed 24 h after electroporation. Displayed is the HCV RNA replication 72 h after treatment. Chimeric prototype SGRs H77 (A&B) or S52 (C&D) harboring the indicated ReEDs and the indicated RAMs were DCV treated as indicated. The dashed grey line marks the 72 h time point for H77/ S52 ΔGDD, replication deficient negative controls. Data are from three independent biological replicates measured in technical duplicates. Each dot depicts the result of one replicate and the bar indicates the mean of all replicates. Statistical significance was determined using a two-sided Student’s t-test. ns = not significant (p > 0.05), * = p < 0.05, ** = p < 0.01.

### Presence of a high replicator ReED is beneficial in sustaining replication upon SOF treatment

Another pilar of anti HCV treatment regimen is the nucleotidic RNA polymerase inhibitor SOF. For this drug, barrier to resistance is quite high thus making SOF RAMs rare. Nevertheless, in the SOF treated patients DAA_6 (gt1b) and DAA_9 (gt3a) the NS5B RAMs C316N [26] and A150V [25] were detected respectively. Both RAMs had a negative effect on RF, although this was more marked for C316N (68% RF reduction) than for A150V (7% RF reduction) (Fig. 7A&B). In both cases, the high replicator ReED induced an increase in RF of over 100-fold, similar to the results for DCV RAMs. Therefore, the high replicator ReED more than compensated for the RAM induced fitness loss. C316N in gt1b and A150V in gt3a appeared to confer no or only very mild treatment resistance when SGRs were treated with SOF at its IC90 (Fig. 7C&D, S3). Still, the results were in line with the observations for DCV with SGRs with a high replicator ReED having a replicative advantage upon DAA treatment.

**Figure 7:**
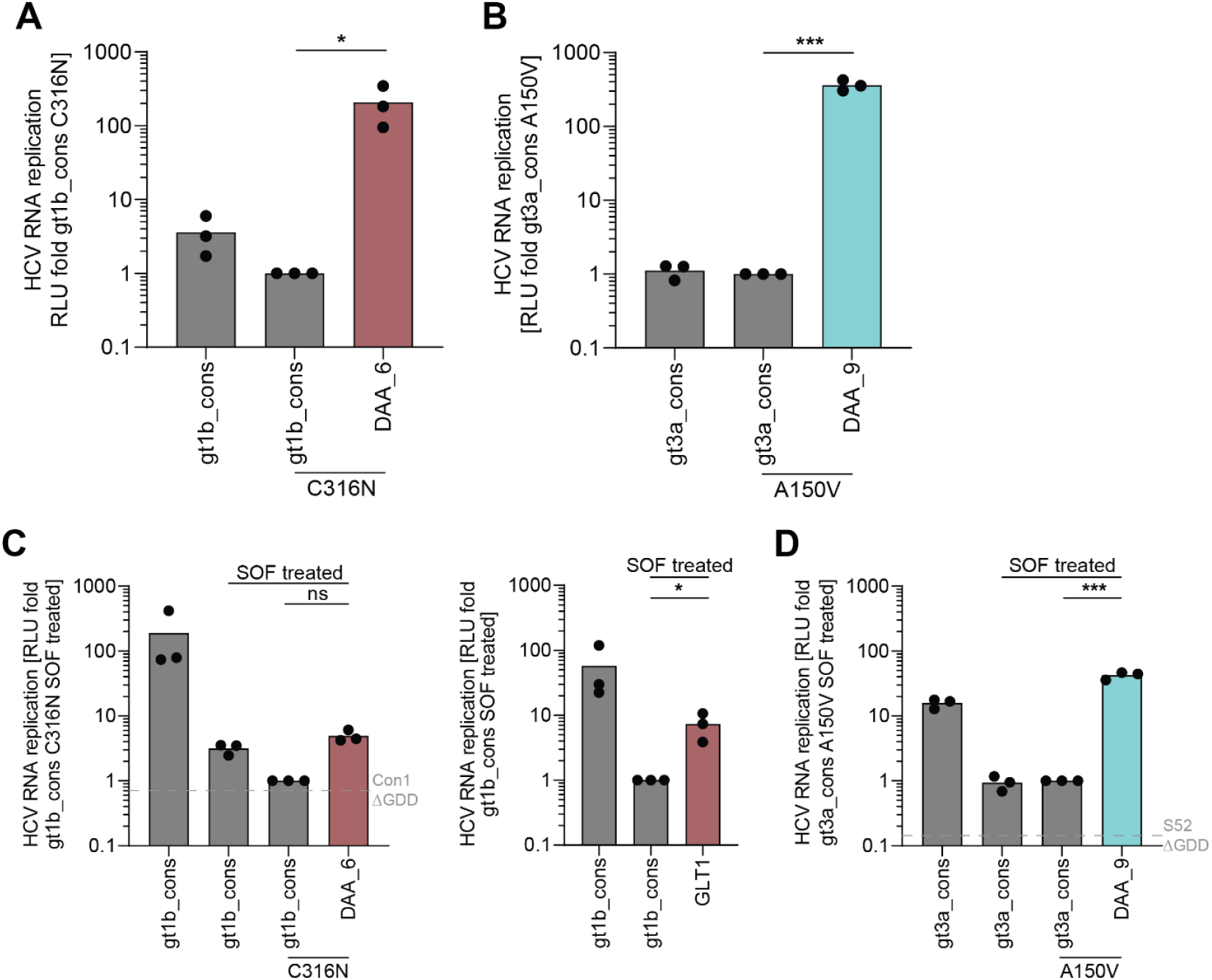
High replicator ReEDs retain an elevated replication level in the presence of SOF RAMs and SOF treatment. Replication studies were performed as described in Fig. 1B. Results were first normalized to 4 h to account for differences in transfection efficiency and then normalized to the RNA replication of the untreated (A&B) or SOF treated (C&D) construct with the gt specific consensus ReED and the indicated RAM. SOF treatment with the IC90 concentration (gt1b 0.77 µM, gt3a 1.23 µM) as determined in Fig. S3 was performed 24 h after electroporation. Displayed is the HCV RNA replication 72 h after electroporation (A&B) or treatment (C&D). Chimeric prototype SGRs Con1 (A&C) or S52 (B&D) harboring the indicated ReEDs and the indicated RAMs were used. The dashed grey line marks the 72 h time point for Con1/ S52 ΔGDD, replication deficient negative controls. Data are from three independent biological replicates measured in technical duplicates. Each dot depicts the result of one replicate and the bar indicates the mean of all replicates. Statistical significance was determined using a two-sided Student’s t-test. ns = not significant (p > 0.05), * = p < 0.05, *** = p < 0.001.

RAM S282T in NS5B was previously shown to confer SOF resistance in cell culture, although at a high fitness cost [25, 27]. This might explain why S282T was not found in our sequencing cohorts and generally is rarely found in patients [11, 12]. Still, S282T is supposed to contribute transiently to treatment failure, we therefore aimed to study its interplay with a high replicator ReED in the context of SOF treatment. We introduced the RAM into S52 containing the DAA_9 ReED since this was already extensively used in the present study and patient DAA_9 failed DAA therapy with a SOF containing regimen. Indeed, S282T induced a 55% fitness reduction in a low replicator SGR harbouring the gt3a consensus ReED and the introduction of the DAA_9 high replicator ReED boosted RF by more than 100-fold (Fig. 8A). The presence of a high replicator ReED did not increase the IC50 values upon SOF treatment (Fig. 8C) but enhanced absolute replication levels by nearly 100-fold in presence of IC90 concentrations (1.23 µM) of SOF compared to the gt3a consensus ReED (Fig. 8B).

**Figure 8:**
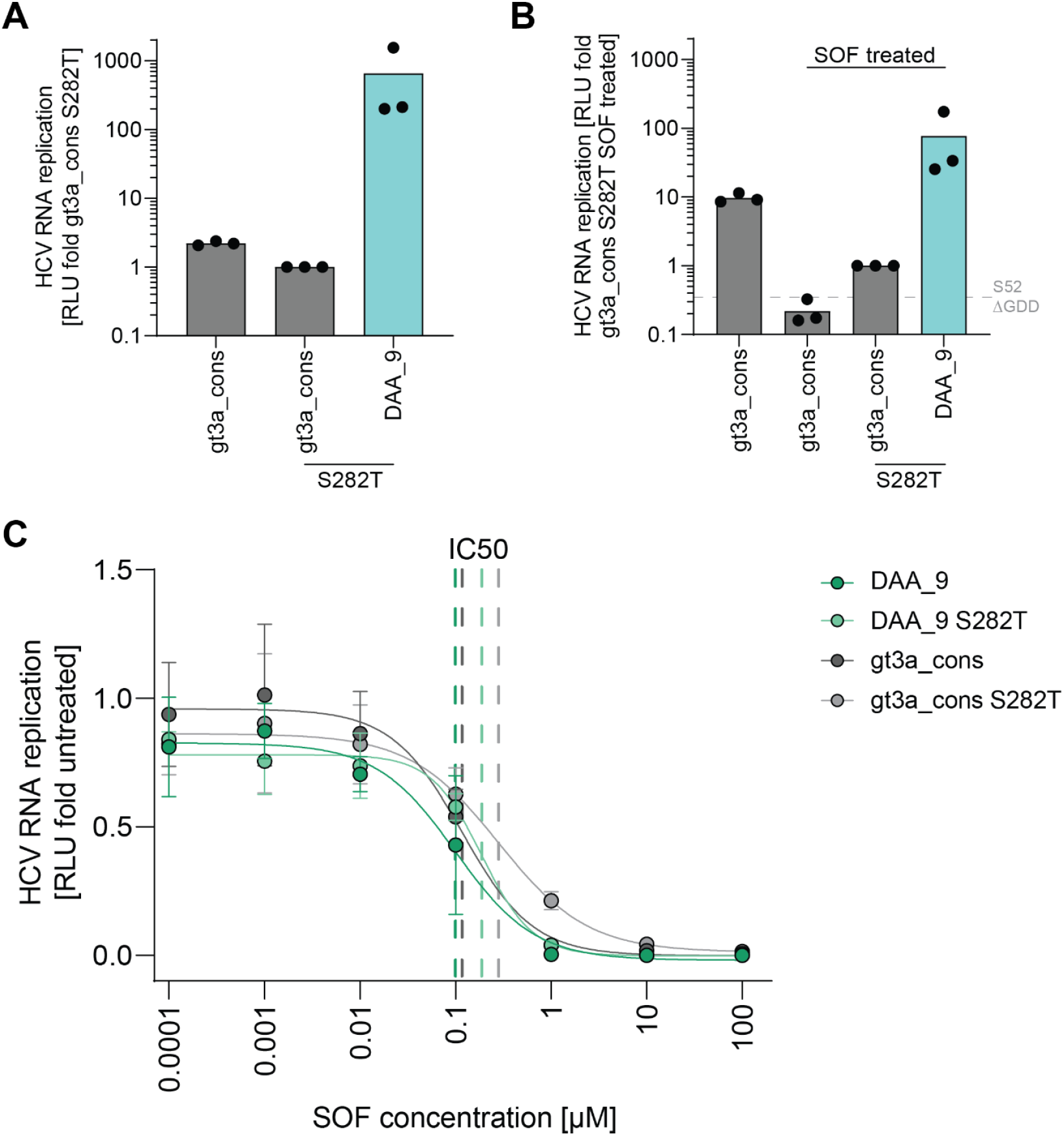
High replicators show superior replication fitness in the context of the SOF RAM S282T. A-C) Replication studies were performed as described in Fig. 1B. Chimeric prototype SGR S52 harboring the indicated ReEDs and the indicated RAM was used. A&B) Results were first normalized to 4 h to account for differences in transfection efficiency and then normalized to the RNA replication of the untreated (A) or SOF treated (B) construct with the gt specific consensus ReED and the indicated RAM. SOF treatment with the IC90 concentration (1.23 µM) was performed 24 h after electroporation. Displayed is the HCV RNA replication 72 h after electroporation (A) or treatment (B). The dashed grey line marks the 72 h time point for S52 ΔGDD, a replication deficient negative control. Data are from three independent biological replicates measured in technical duplicates. Each dot depicts the result of one replicate and the bar indicates the mean of all replicates. C) Electroporated cells were treated with variable SOF concentrations 24 h after electroporation. Depicted is the replication 72 h after treatment. Data are from three independent biological replicates measured in technical duplicates with each dot depicting the result of one replicate. IC50 was determined through a nonlinear fit shown as a continuous line, the dashed line marks the IC50.

Overall, the SOF data are in line with the DCV data highlighting that a high replicator ReED can benefit HCV during DAA treatment not by per se reducing the drug’s direct antiviral effect but by creating a replicative buffer, helping the virus to maintain higher replication levels. Additionally, the ReED’s replication enhancing effect mitigated the fitness loss induced by RAMs such as S282T facilitating their emergence. Thus, the synergy of increased RF mediated by ReED mutations and RAMs might contribute to DAA treatment failure implying that patients failing DAA therapy should not only be checked for the presence of RAMs but also for mutations in the ReED.

## Discussion

In this study we addressed the impact of HCV replication fitness on therapy outcome of IFN- and DAA-based treatment regimens. Our data show that high viral replication fitness supports successful IFN treatment but can contribute to DAA therapy failure.

The correlation between an accumulation of ISDR mutations and improved IFN therapy response was initially identified in Japanese patients [18, 28] with meta analyses of international follow-up studies confirming this connection for gt1b patients in general [29-31]. We now phenotypically analysed a panel of viral sequences from the initial study [18], demonstrating that all patients infected with a high replicator variant indeed were successfully treated with IFN. However, some successfully cured patients harboured low replicators, demonstrating that high replication capacity supported IFN treatment, but was not a prerequisite for response. Our results are in line with one previous study, where the ISDR of P1_IFN(s) was introduced into the RpN and RpJG gt1b SGRs stimulating RF [32]. However, this and other previous studies [33, 34] on the impact of ISDR mutations on IFN sensitivity in cell culture were limited by the use of cell culture adaptive mutations, masking the replication enhancing effect of the ReED.

The reason for the increased IFN sensitivity of high replicators remains a matter of speculation. While NS5A was shown to counteract the IFN response, no differences induced by ISDR mutations could be observed in previous studies [35, 36] in line with our results revealing similar IC50 values for IFN treated low and high replicators. Nevertheless, RF of high replicators remained elevated in the presence of given concentrations of IFN. It is therefore tempting to speculate that increased antigen levels might lead to a stronger stimulation of the adaptive immune system, facilitating detection and elimination of infected cells. More refined cell culture systems also including components of the adaptive immune system [37, 38] would be needed to clarify this hypothesis. However, impairment of antigen presentation was also recently suggested as the cause of IFN treatment failure in patients harbouring a functional IFN lambda 4 gene [39], a genetic background that was also associated with a higher prevalence of high replicators [17]. Impaired antigen presentation in such patients might further limit the stimulation of adaptive immune responses by high replicator variants, thereby masking their contribution to treatment success in human populations with high abundance of the unfavourable IFN lambda alleles. The complex contribution of ISDR mutations and human genetics to IFN treatment outcome might further explain, why studies for other HCV gts rarely identified connections between ISDR mutations and IFN sensitivity [40-42], although we found that ISDR mutations can increase RF in all major HCV gts [17]. Indeed, this study shows that gt1a isolates 1a_IFN_1 and 1a_IFN_2 from patients with a good IFN treatment response [19], both were high replicator variants, implying a connection between the two parameters also beyond gt1b.

Furthermore, we were able to show that high replicator ReEDs can be found in some DAA failure patients with the high replicator phenotype remaining observable in the presence of RAMs. Upon DAA treatment, the high replicators were able to sustain higher levels of replication thus potentially aiding the virus population in withstanding antiviral treatment. Nevertheless, RF did not affect the SOF IC50 for gt3a isolates which is in line with our previous findings for DCV and SOF treatment in gt1b [17]. Also, for cell culture adapted gt2a variants, a connection between higher RF and the ability to sustain infectivity upon DAA treatment was shown [43, 44].

Our cohort gave us the opportunity to work with authentic combinations of RAMs and ReEDs from treatment failure patients. Since the cohort mainly consisted of patients treated with first generation NS5A inhibitors and SOF, we focused on these drugs and their associated RAMs. In NS5A, L31M and Y93C in gt1a and Y93H in gt1b showed similar effects on RF and drug resistance as previously described [45]. For gt3a, the lack of a reduction in RF induced by Y93H observed by us was surprising, since a fitness cost of this mutation was described several times although with varying magnitudes [22, 46, 47], probably due to the use of different cell culture models. The SOF RAMs A150V and C316N have previously been associated with treatment failure in patients, but in line with our data, showed only moderate or no impact on replication fitness and drug response in cell culture [25, 26, 48]. In contrast, S282T caused a strong reduction in RF, in line with literature [25, 27]. In general, it has been shown that secondary mutations could compensate for the fitness loss of some RAMs [22, 49], a concept of epistasis that is supported by modelling approaches [50]. However, previous studies were not able to identify mutations compensating for the S282T induced fitness loss in cell culture [51], but associations between mutations in NS2 and NS3 and SOF treatment failure [52, 53]. In our study, the ReED actually enhanced RF to a much larger extent than the RAM induced fitness loss, resulting upon DAA treatment in replication levels comparable to untreated low replicator isolates. Still, it remains challenging to translate the drug concentrations used in vitro to an in vivo setting. One study reported that in the explanted liver of transplantation patients, the SOF concentrations ranged from 0.382 µM to 252 µM with a median of 70.3 µM [54]. So, at least for some patients the 1.23 µM SOF concentration used here for gt3a constructs might be comparable to the situation in vivo, a condition under which a high replicator with S282T still showed replication over 100-fold higher than a replication deficient control.

In general, it is paradox that the same phenotype, high replication fitness, can enhance treatment success for antiviral IFN therapy but reduce it for DAA treatment. This highlights the relevance of a separation between intrinsic fitness determined without external factors like drug treatments or immune pressure and effective fitness in the presence of these factors [55, 56]. IFN likely acts indirectly on the virus thus allowing a high replicator to produce high levels of antigen, subsequently enhancing the antiviral immune response leading to HCV elimination. Therefore, the highest effective fitness allowing for persistence is achieved by low replicators. DAAs directly inhibit viral functions, eliminating the virus by preventing any viral antigen production in a largely immune system independent manner. Here, highest effective fitness allowing for persistence is achieved by high replicators preserving replication also by compensating RAM induced fitness costs.

Furthermore, the antiviral treatment causes a very sudden shift in the effective fitness of HCV variants impacting viral evolution. In general, the ISDR sequence was shown to be quite stable over time in chronically infected patients [57, 58]. Nevertheless, upon IFN treatment it can change rapidly as it was previously shown that species with ISDR mutations vanished upon IFN treatment while species with less ISDR mutations persisted arguing for a selection for low replicators [18, 59]. Along these lines, we identified two patients where a high replicator dominated only after failed DAA treatment, further indicating that a selection for high RF upon DAA treatment is beneficial for the virus. Similarly, RAMs mostly emerge upon treatment since only then they offer a fitness advantage. After waning of DAAs, the fitness advantage of RAMs like S282T vanishes often leading to a reversion back to wildtype [60] which can happen within days in the case of S282T [11, 61]. Along these lines, high replicator ReEDs appear to be unfavourable in immunocompetent patients [17], so the ReED might revert to a low replicator version after treatment either through reverting to a more wildtype-like sequence or by acquiring replication reducing mutations in the C-term of the ReED as observed for P12_IFN(s). Thus, some sequencing studies might sample too late after treatment failure for the detection of a high replicator ReED. Furthermore, more serial samples would be required to assess what comes first: replication enhancing mutations in the ReED or RAMs. Currently, sequencing of HCV in patients who failed DAA therapy usually focusses on known hotspots for RAMs like domain 1 of NS5A [62-64]. Our results indicate that extending the sequenced region of NS5A to also include the ReED would improve the understanding of reasons for treatment failure in individual patients and might help to support decisions for alternative treatment regimens.

In conclusion, our study reveals how the differences in the way antiviral therapies act also shape the viral resistance mechanisms. In IFN treatment which likely suppresses the virus through immune mediated effects, high genome replication fitness appeared to be detrimental for HCV, likely by supporting adaptive immunity. For DAA treatment, our data indicate that elevated replication fitness might help the virus to persist in presence of antiviral drugs directly by preserving higher replication levels and indirectly by compensating the RAMs’ fitness costs facilitating their establishment. Thus, factors determining the outcome of current HCV therapies go beyond RAMs and our data argue for monitoring of sequence signatures of enhanced replication fitness in patients failing DAA therapy.

## Methods

### Ethics statement

This study was approved by the ethics committee of the medical faculty of Heidelberg University (ethics votes: S-743/2023). The studies were conducted in accordance with the local legislation and institutional requirements. The participants provided their written informed consent to participate in this study. The usage of individuals’ blood samples and the retrospective collection of limited pseudonymized patient data were approved by the ethics committee of the University Hospital Frankfurt, Germany (ethics vote: 16/15).

### Patients

The IFN treatment cohort [18], patients 1a_IFN_1 & 1a_IFN_2 [19], the HCV Research UK cohort [20] and the PEPSI cohort were published previously [21]. Gt1b samples received from the European DAA resistance database at the University Hospital Frankfurt, Germany, were selected to be either DAA treatment failure without detected RAM (n = 10), or because of the presence of either Y93H (n = 15) or L31M+Y93H (n = 13) RAMs in NS5A. Details about the DAA treatment of the patients whose ReEDs were characterised in cell culture can be found in Table S3.

### Sequence analysis

Sequences were analysed as previously described [17]. RAMs were determined using HCV GLUE [65].

### HCV replication in cell culture

Cloning of plasmids encoding HCV SGRs, in vitro transcription, culturing of Huh7-Lunet SEC14L2 cells, transfection of these cells and luciferase activity measurements were previously described in detail [17].

### Drug treatment

Cells transfected with HCV IVT RNA through electroporation were treated with IFN2α (Active Bioscience), DCV (Bristol Myers Squibb) or SOF (MedChemExpress) 24 h after electroporation. Samples for luciferase assays were harvested 72 h after treatment. IC50 and IC90 values were determined employing a nonlinear fit in GraphPad Prism (Graphpad Software).

### Statistical analysis

Statistical analyses were performed using GraphPad Prism (Graphpad Software) or R. For numerical data, a two-sided Student’s t-test was applied when normal distribution was assumed.

## Acknowledgements

We thank R. Klein and U. Herian for excellent technical assistance. For the generous gift of plasmids, we are grateful to J. Bukh (H77) and C. Rice (S52). The authors wish to acknowledge the role of HCV Research UK (funded by the Medical Research Foundation, award number C0365) in collecting and making available the data used in the generation of this publication.

## Data availability

ReED sequence data of 38 DAA failure patients from the European DAA resistance database at the University Hospital Frankfurt, Germany are deposited in GenBank under accessions PX738258-PX738295. All other data generated during this study are included in this article and its supplementary files.

## Author contributions

HGTS: Investigation, Visualization, Writing - Original Draft, Writing - Review & Editing; TA: Investigation, Visualization, Writing - Original Draft, Writing - Review & Editing; CF: Investigation, Writing - Review & Editing; MB: Investigation, Writing - Review & Editing; EH: Investigation, Writing - Review & Editing; JM: Resources, Writing - Review & Editing; CS: Resources, Writing - Review & Editing; JD: Resources, Writing - Review & Editing; RK: Resources, Writing - Review & Editing; VL: Conceptualization, Supervision, Funding acquisition, Writing - Original Draft, Writing - Review & Editing; PR: Conceptualization, Software, Investigation, Visualization, Supervision, Writing - Original Draft, Writing - Review & Editing

