## Supplementary material for "Hepatitis C virus replication fitness as a determinant of antiviral therapy outcome": Table S3

Supplementary Figures S1-S3

Supplementary Tables S1-S3

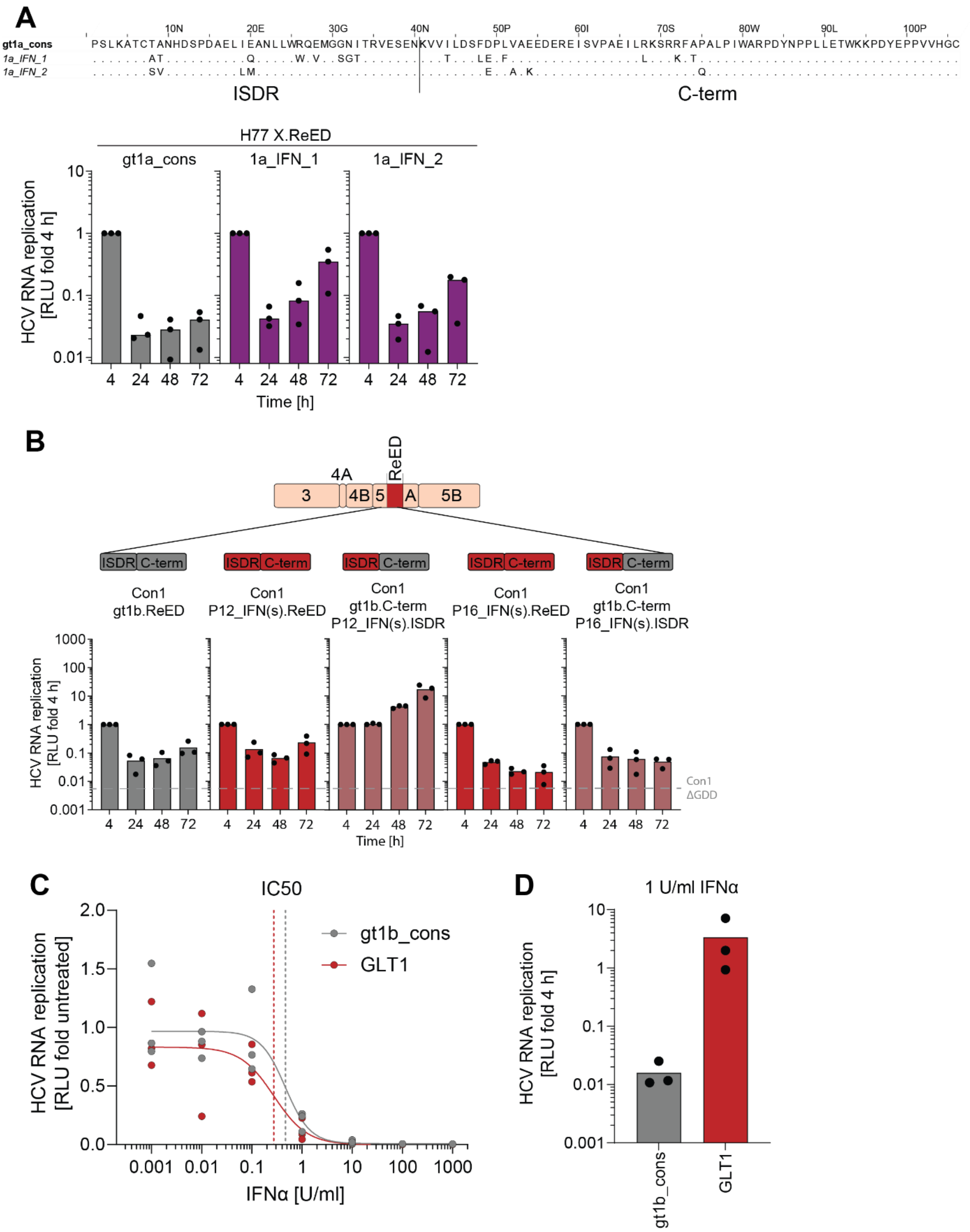


**Figure S1: Determinants of IFN sensitivity in gt1a and gt1b.** A) Alignment of the ReED amino acid sequences of 1a_IFN_1 and 1a_IFN_2 [19] with a genotype 1a consensus ReED serving as a reference. Dots indicate that an amino acid is identical to the consensus (upper panel). Replication studies were performed as described in Fig. 1B using H77 based SGRs harbouring the indicated ReED (lower panel). Data are from three independent biological replicates measured in technical duplicates. Each dot depicts the result of one replicate and the bar indicates the mean of all replicates. B) Con1 based SGRs harboring either the complete ReED from isolates P12_IFN(s)/P16_IFN(s) or just the ISDR of these isolates combined with the ReED C-term of the gt1b consensus were used for replication studies as described in Fig. 1B. The dashed grey line indicates the 72 h time point for Con1 ΔGDD, a replication deficient negative control. Data are from three independent biological replicates measured in technical duplicates. Each dot depicts the result of one replicate and the bar indicates the mean of all replicates. C) Huh7-Lunet SEC14L2 cells electroporated with Con1 SGRs harboring the indicated ReED were treated with variable IFN concentrations 24 h after electroporation. Depicted is the replication 72 h after treatment. Data are from three independent biological replicates measured in technical duplicates with each dot depicting the result of one replicate. IC50 was determined through a nonlinear fit shown as a continuous line, the dashed line marks the IC50. D) Data from panel C showing the replication 72 h after treatment with 1 U/ml IFN normalized to the input 4 h after electroporation.

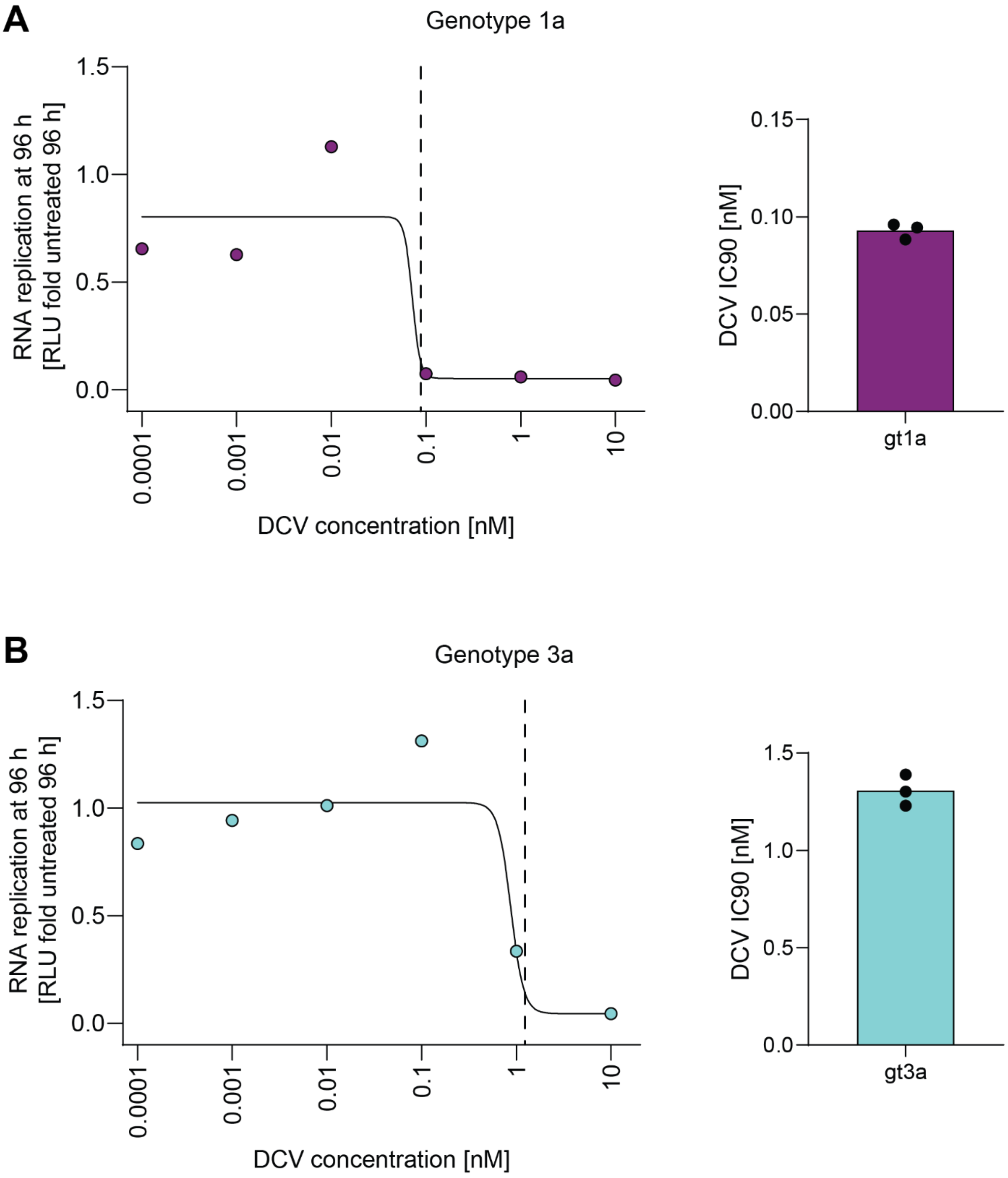

**Figure S2: Determination of DCV IC90s.** Replication studies were performed as described in Fig. 1B. Cells were treated with variable DCV concentrations 24 h after electroporation. The SGRs H77 gt1a_cons.ReED (A) and S52 gt3a_cons.ReED (B) were used. Depicted is the replication 72 h after treatment of one representative experiment (left panels) out of a total of three independent biological replicates measured in technical duplicates. The IC90 (dashed line) was determined through a nonlinear fit shown as a continuous line. In the right panels, the IC90s of each replicate are shown as dots with the bar representing the mean IC90 used for subsequent experiments: gt1a 0.09 nM, gt3a 1.3 nM.

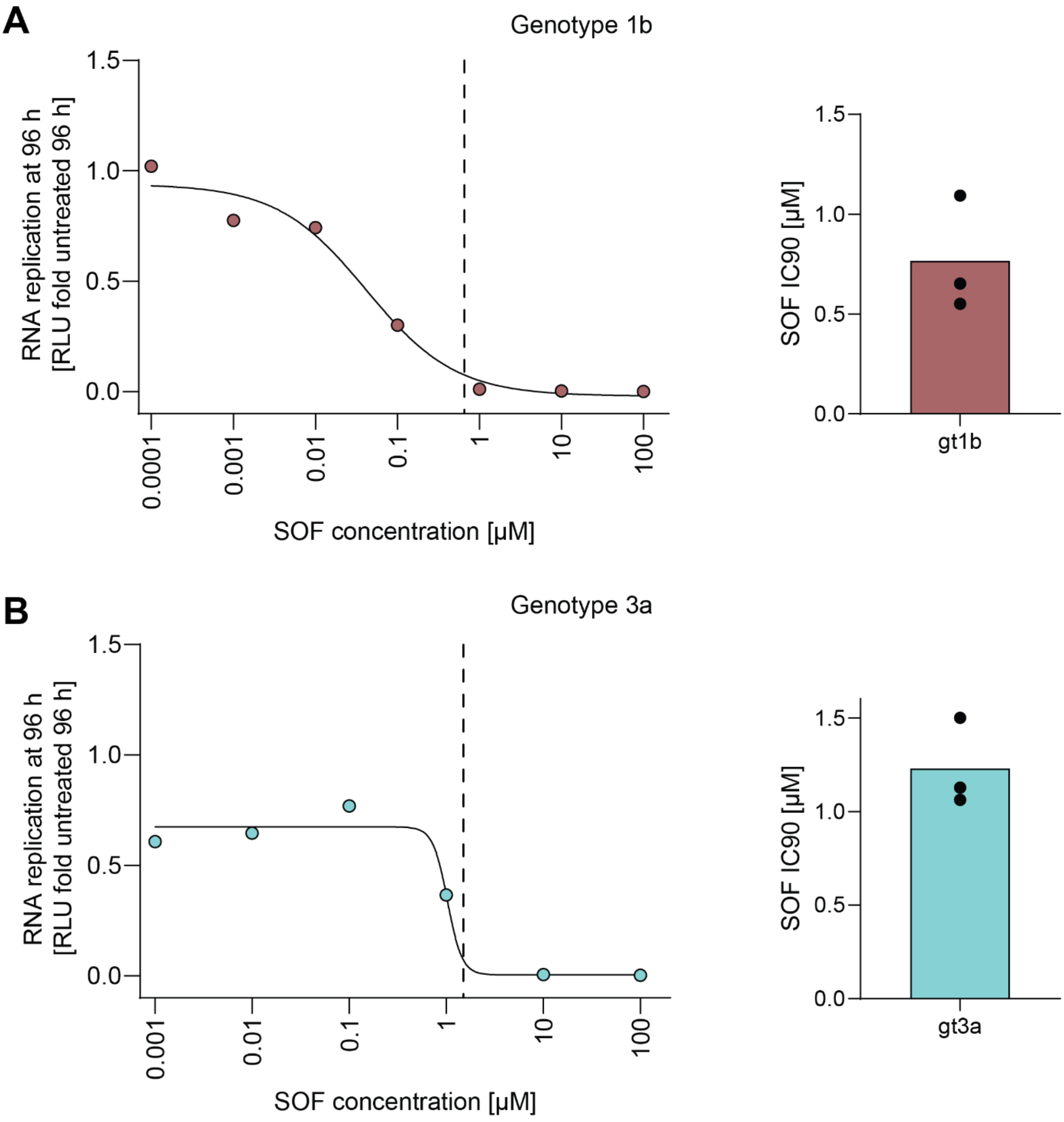


**Figure S3: Determination of SOF IC90.** Replication studies were performed as described in Fig. 1B. Cells were treated with variable SOF concentrations 24 h after electroporation. The SGRs Con1 gt1b_cons.ReED (A) and S52 gt3a_cons.ReED (B) were used. Depicted is the replication 72 h after treatment of one representative experiment (left panels) out of a total of three independent biological replicates measured in technical duplicates. The IC90 (dashed line) was determined through a nonlinear fit shown as a continuous line. In the right panels, the IC90s of each replicate are shown as dots with the bar representing the mean IC90 used for subsequent experiments: gt1b 0.77 µM, gt3a 1.23 µM.

**Table S1: DNA plasmids used and generated during this project.**

| Name | Description | GenBank accession |
|---|---|---|
| pFK i341 NS3-3' Con1 | Bicistronic SGR of the prototype gt1b wildtype isolate | CAB46677 |
| pFK i341 NS3-3' Con1 gt1b_cons.ReED | Con1 based chimera harboring the gt1b consensus ReED | |
| pFK i341 NS3-3' Con1 GLT1.ReED | Con1 based chimera harboring patient derived ReEDs | OM222702 |
| pFK i341 NS3-3' Con1 P1_IFN(s).ReED | | |
| pFK i341 NS3-3' Con1 P2_IFN(s).ReED | | |
| pFK i341 NS3-3' Con1 P3_IFN(s).ReED | | |
| pFK i341 NS3-3' Con1 P10_IFN(s).ReED | | |
| pFK i341 NS3-3' Con1 P11_IFN(s).ReED | | |
| pFK i341 NS3-3' Con1 P12_IFN(s).ReED | | |
| pFK i341 NS3-3' Con1 P13_IFN(s).ReED | | |
| pFK i341 NS3-3' Con1 P14_IFN(s).ReED | | |
| pFK i341 NS3-3' Con1 P15_IFN(s).ReED | | |
| pFK i341 NS3-3' Con1 P16_IFN(s).ReED | | |
| pFK i341 NS3-3' Con1 P17_IFN(s).ReED | | |
| pFK i341 NS3-3' Con1 P4_IFN(r).ReED | | |
| pFK i341 NS3-3' Con1 P7_IFN(r).ReED | | |
| pFK i341 NS3-3' Con1 P8_IFN(r).ReED | | |
| pFK i341 NS3-3' Con1 DAA_5.ReED | | PX738278 |
| pFK i341 NS3-3' Con1 DAA_6.ReED | | |
| pFK i341 NS3-3' Con1 gt1b_cons.ReED Y93H | Con1 based chimera harboring patient derived ReEDs and RAMs | |
| pFK i341 NS3-3' Con1 GLT1.ReED Y93H | | |
| pFK i341 NS3-3' Con1 DAA_5.ReED Y93H | | |
| pFK i341 NS3-3' Con1 DAA_6.ReED Y93H | | |
| pFK i341 NS3-3' Con1 gt1b_cons.ReED C316N | | |
| pFK i341 NS3-3' Con1 DAA_6.ReED C316N | | |
| pFK i341 NS3-3' Con1 gt1b.C-term P12_IFN(s).ISDR | Con1 based chimera harboring the gt1b consensus C-term and patient derived ISDRs | |
| pFK i341 NS3-3' Con1 gt1b.C-term P16_IFN(s).ISDR | | |

| | | |
|---|---|---|
| pFK i341 NS3-3' H77 | Bicistronic SGR of the prototype gt1a wildtype isolate | AF009606 |
| pFK i341 NS3-3' H77 gt1a_cons.ReED | H77 based chimera harboring the gt1a consensus ReED | |
| pFK i341 NS3-3' H77 IFN_1.ReED | H77 based chimera harboring patient derived ReEDs | EF407414 |
| pFK i341 NS3-3' H77 IFN_2.ReED | | EF407416 |
| pFK i341 NS3-3' H77 DAA_1.ReED | | |
| pFK i341 NS3-3' H77 DAA_2.ReED | | |
| pFK i341 NS3-3' H77 DAA_3.ReED | | |
| pFK i341 NS3-3' H77 DAA_4.ReED | | |
| pFK i341 NS3-3' H77 FCH1a_2.ReED | | JQ914274 |
| pFK i341 NS3-3' H77 gt1a_cons.ReED L31M | H77 based chimera harboring patient derived ReEDs and RAMs | |
| pFK i341 NS3-3' H77 DAA_2.ReED L31M | | |
| pFK i341 NS3-3' H77 gt1a_cons.ReED Y93C | | |
| pFK i341 NS3-3' H77 DAA_3.ReED Y93C | | |
| pUC i387 NS3-3' S52 | Bicistronic SGR of the prototype gt1b wildtype isolate | GU814263 |
| pUC i387 NS3-3' S52 gt3a_cons.ReED | S52 based chimera harboring the gt3a consensus ReED | |
| pUC i387 NS3-3' S52 DAA_7.ReED | S52 based chimera harboring patient derived ReEDs | |
| pUC i387 NS3-3' S52 DAA_8.ReED | | |
| pUC i387 NS3-3' S52 DAA_9.ReED | | |
| pUC i387 NS3-3' S52 gt3a_cons.ReED Y93H | S52 based chimera harboring patient derived ReEDs and RAMs | |
| pUC i387 NS3-3' S52 DAA_9.ReED Y93H | | |
| pUC i387 NS3-3' S52 gt3a_cons.ReED A150V | | |
| pUC i387 NS3-3' S52 DAA_9.ReED A150V | | |
| pUC i387 NS3-3' S52 gt3a_cons.ReED Y93H A150V | | |
| pUC i387 NS3-3' S52 DAA_9.ReED Y93H A150V | | |
| pUC i387 NS3-3' S52 gt3a_cons.ReED S282T | | |
| pUC i387 NS3-3' S52 DAA_9.ReED S282T | | |

**Table S2: Oligo nucleotides used for ReED amplification from patient sera.**

| Name | Sequence (5‘-3‘) | Description | Gt |
|---|---|---|---|
| A_9416 | CAGGATGGCCTATTGGCCTGGAG | cDNA generation | 1b |
| S_6767 | ATTCCAGGTCGGGCTCAA | Nested PCR from cDNA to amplify the ReED | 1b |
| S_6843 | ACTTCCATGCTCACCGACCC | | 1b |
| A_7501 | AGAGGGGGCATGGAGGAGTA | | 1b |
| A_7552 | ACGGTAGACCAAGACCCGTC | | 1b |

**Table S3: Characteristics of DAA failure patients whose ReED was analysed in cell culture.** na = not available.

| Name | Source | Start date of unsuccessful treatment | Drug scheme of unsuccessful treatment | Notes |
|---|---|---|---|---|
| DAA_1 | HCV Research UK | 10.2014 | Sofosbuvir, Daclatasvir, Ribavirin | Responder-Relapser |
| DAA_2 | PEPSI | 04.2015 | Ledipasvir, Sofosbuvir | Responder-Relapser |
| DAA_3 | PEPSI | 05.2016 | Ledipasvir, Sofosbuvir | Responder-Relapser |
| DAA_4 | PEPSI | 07.2016 | Ledipasvir, Sofosbuvir | Responder-Relapser |
| DAA_5 | European DAA resistance database | na | Ledipasvir, Sofosbuvir, Ribavirin | Responder-Relapser |
| DAA_6 | PEPSI | 04.2015 | Ledipasvir, Sofosbuvir, Ribavirin | Responder-Relapser (First treatment) |
| | | 03.2016 | Simeprevir, Sofosbuvir, Ribavirin | Responder-Relapser (Second treatment) |
| DAA_7 | HCV Research UK | 08.2014 | Sofosbuvir, Daclatasvir, Ribavirin | Responder-Relapser |
| DAA_8 | HCV Research UK | 07.2014 | Sofosbuvir, Daclatasvir, Ribavirin | Responder-Relapser |
| DAA_9 | PEPSI | 09.2015 | Daclatasvir, Sofosbuvir | Virological failure |
| 3a_LTX_2 | HCV Research UK | 09.2014 | Sofosbuvir, Daclatasvir, Ribavirin | Responder-Relapser, LTX in 03.2011 |
